# Electrochemical biosensors based on peptide-kinase interactions at the kinase docking site

**DOI:** 10.1101/2021.11.16.468793

**Authors:** Pralhad Namdev Joshi, Evgeniy Mervinetsky, Ohad Solomon, Yu-Ju Chen, Shlomo Yitzchaik, Assaf Friedler

## Abstract

Kinases are important cancer biomarkers and are conventionally detected based on their catalytic activity. Kinases regulate cellular activities by phosphorylation of motif-specific multiple substrate proteins, resulting in lack of selectivity of activity-based kinase biosensors. We present an alternative approach of sensing kinases based on the interactions of their allosteric docking sites with a specific partner protein. The new approach was demonstrated for the ERK2 kinase and its substrate ELK-1. A peptide derived from ELK-1 was bound to a gold electrode and ERK2 sensing was performed by electrochemical impedance spectroscopy. The sensors showed high level of target selectivity for ERK2 when compared with p38γ kinase and BSA. ERK2 was detected in its cellular concentration range, 0.2-8.0 μM. Using the flexibility of peptide design, our method is generic for developing sensitive and substrate-specific biosensors and other disease-related enzymes based on their interactions.

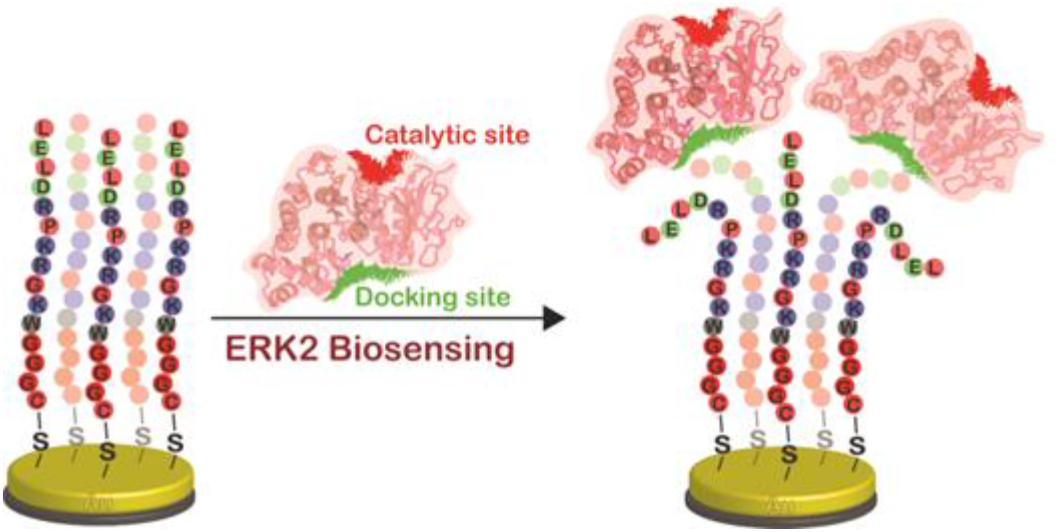

## 1. Introduction

Kinases plays critical roles in regulating signal transduction and various cellular processes. Aberrant overexpression of a kinase could dynamically phosphorylate its various substrates to ubiquitously influence multiple signal transduction pathways and causes initiation and progression of diseases.^1^ Therefore, kinases are important cancer biomarkers since they are overexpressed in many tumors.^2^ The development of highly sensitive biosensors for cancer biomarkers is a key goal in the early diagnosis of cancer^3^ Mass spectrometry-based phosphoproteomics techniques are currently used to identify disease-associated alteration of kinases and their substrates for biomarker discovery.^4^ Although MS can identify the kinase-substrate pair as biomarker candidates, it requires complicated protocols and analytical tool, which limit analytical sensisitivity and rubustness. Current techniques for detecting proteins include enzyme-linked immunosorbent assays (ELISA) as one of the most common method.^5^ ELISA is developed based on antibodies for their recognition of a specific protein or the target biomolecules. However, the ELISA assay relies on the availablility and quality of priminary antibody and secondary antibody labed on reporter enzymes,^6^ which are not easily obtained. Interaction specificity of antibody-antigen was also utilized for the biosensors discovery studies.^7^ Optical sensors such as fluorescent probes and FRET-based sensors use fluorescence or chemiluminescence for the identification of the biological target.^8^ However, these diagnosis approaches rely on reporter molecules, fluorescently labeled or radiolabeled biomolecules, and various chromatographic methods for biomolecule separation from the complex biological samples. In addition, the attached probe may alter the specificity of the interaction between the biomolecules.^9^ The low stability of the antibodies limits their utility and thus the shelf-life of antibody-based biosensors, since antibodies may undergo structural changes and lose their recognition capacity if kept at the wrong conditions.^10^ In additiion, the difficulty in obtaining high specificity to the f target protiens in complex samples is the major factor that casues non-specific detection and limits using antibdy-based assays.

Electrochemical biosensors are important tools in bio-analytics, kinase detection, probing natural systems, cancer diagnosis, preparation of peptide-based biosensors, and foster novel kinase monitors.^11^ Peptide-based electrochemical biosensors^12^ use peptides bound to electrodes that are used for detecting an analyte protein, based on their interaction. The electrochemical response is measured to correlate with the concentration of a peptidic probe from the complex of probe-kinase. Electrochemical impedance spectroscopy (EIS) is an excellent tool for electrochemical biosensing. It can measure the change in the impedance signal due to complex formation between the peptide-based sensor and the analyte protein.^13^ The EIS technique is label-free and does not require additional reagent use.^14^ Recently, we developed electrochemical peptide-based biosensors that sense kinases based on their enzymatic activity.^15^ For example, we developed a peptide-based biosensor for the ERK2 kinase, which is overexpressed in non-small-cell lung cancer (NSCLC).^16^ A peptide was designed based on overly activacted phosphorylation from the HDGF protein, a substrate protein phosphorylated by ERK2^17^, in tumors among NSCLC patients.^16^ The HDGF peptide, derived from its phosphorylation site, was bound to the electrode surface and the electrochemical signal resulting from the phosphorylation of peptide on the electrode surface was used for identifying the target kinase. Another biosensing approach for enzymes, shown in our previous work, described the effects of molecular dipoles on semiconducting interfaces, showing the molecular recognition is independent from the biocatalytic activity of acetylcholine esterase.^18^

Despite the advantages of sensing enzymes based on their catalytic activity, the approach is limited and improved analytical methods should be developed first, many protein disease biomarkers are not enzymes and thus do not have catalytic activity to provide reporter signals. Moreover, specific interactions with the substrate-specific peptide probe can result in more specific sensing compared to relying solely on the catalytic activity, whose specificity is lower. Most importantly, given the different phosphorylation-mediated functions that range between tens and hundreds of subtsrate proteins for the same kinase, a substrate-specific peptide assay provides better disease specificity. To address this, we describe peptide-based electrochemical biosensors that sense kinases based on their docking site interactions instead of their catalytic activity. To achieve this, we targeted the docking of ERK2 and not its catalytic site.^19^

ERK2 contains several major domains: the activation loop, the ATP-binding site and the active site. In addition it contains two docking sites that bind ERK2 partner proteins: (i) the sited-recruitment site (DRS), which consists of the negatively charged D316 and D319 and the hydrophobic H123 to L155 that is far from the catalytic site and binds specifically to the docking motifs in its target proteins (Fig. 1).^20^; (ii) The F-site recruitment site (FRS), which consists of residues Leu-198, Tyr-231, Leu-232, Leu-235, and Tyr-261 on ERK2.^21^ The two docking domains confer the specificity on ERK2 to its target proteins. The charged and hydrophobic residues from the ELK-1 bind to the DRS and shows selectivity towards it so we targeted the DRS and not FRS.

**Fig. 1.**
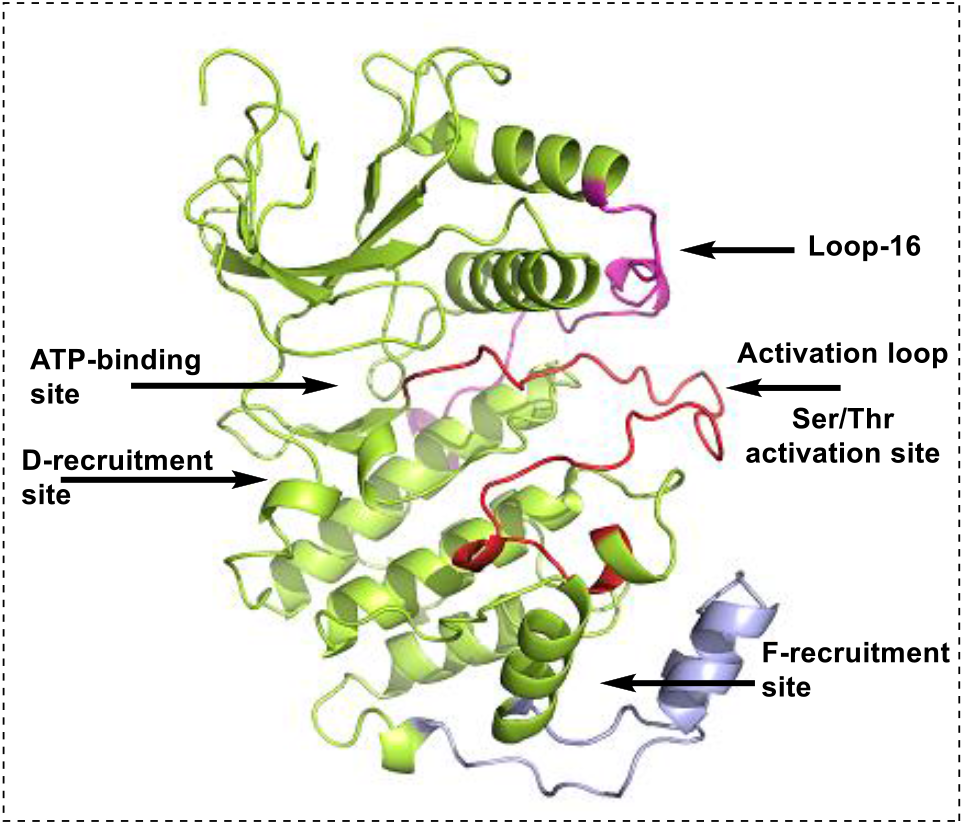
Structure of ERK2. Shown are the different domains - The D-recruitment site bearing the common-docking domain; the ATP-binding site; the activation loop that has the activation site for Ser/Thr phosphorylation; the F-recruitment site, which is an additional interaction site for the substrate protein.

Here we describe the development of a sensor based on the interaction of ERK2 with a peptide derived from the docking motif in the D domain of its binding partner protein ELK-1.^22^ The peptide sequence contains hydrophobic and charged residues (residues 311-327 in ELK, QKGRKPRDLELPLSPSL).^23^ The peptide sequence from a D domain binds the DRS on ERK2 (Fig. 2). The sensor is highly sensitive and selective and is a proof of concept for sensing kinases not based on their activity but rather on specific interactions with their substrate proteins.

**Fig. 2.**
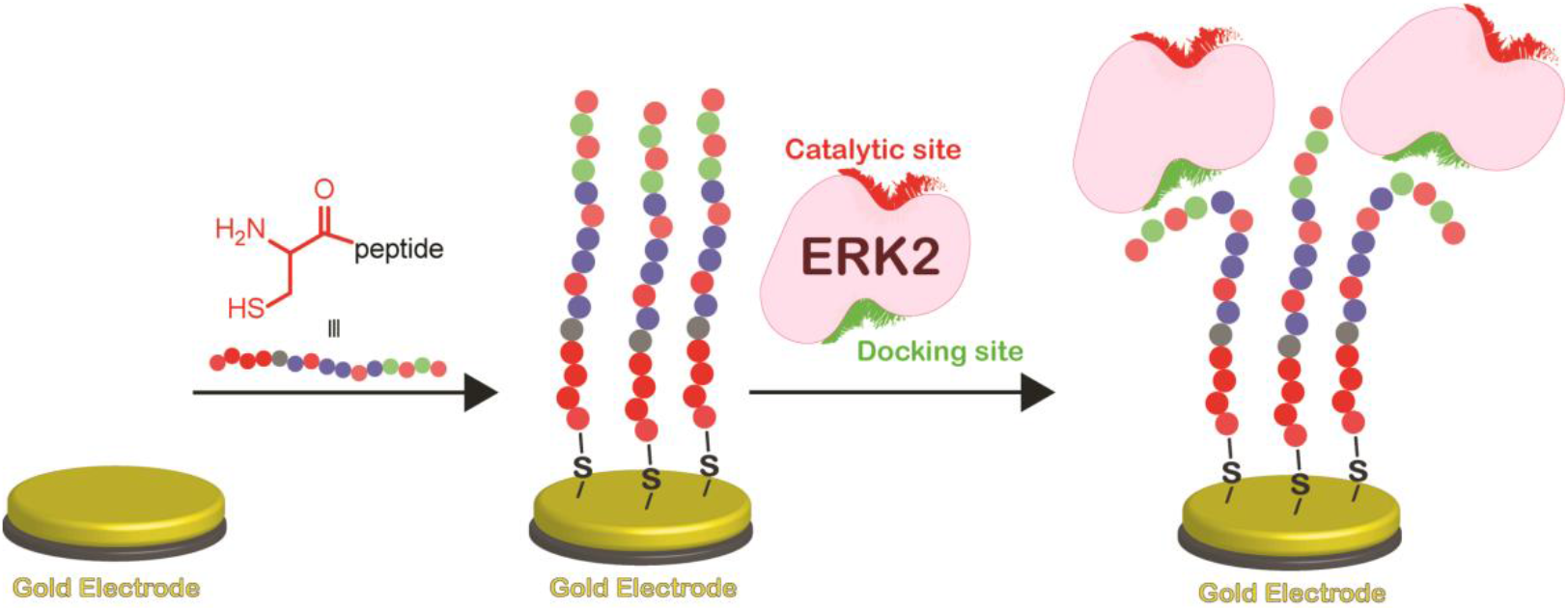
Interaction based peptide-kinase sensor detecting the enzyme based on docking site and not catalytic site.

## 2. Experimental

### 2.1 Peptide synthesis, labeling and purification protocol

Peptides were synthesized using standard Fmoc chemistry using N,N′-Diisopropylcarbodiimide (DIC) and Oxyma as coupling reagents. The synthesis was carried on an automated Liberty Blue Microwave-Assisted Peptide Synthesizer (CEM). A Trp residue was added to the ELK-1_312-321_ peptides to determine their concentrations using UV/Vis spectroscopy. For fluorescence anisotropy binding studies the peptide (2) was synthesized with 5(6)-carboxyfluorescein at its N-terminus as described (Fig. S1).^24^ The peptides were purified on preparative-HPLC (Waters 150 Q LC system using an XSelect C18 column (5 μm, 30 × 250 mm).) Linear gradients from 5 to 60% of MeCN (with 0.1% TFA, buffer B) in water (with 0.1% TFA, buffer A) over a 42 minutes were used to elute the peptides. The purity was analyzed using an analytical HPLC (Merck-Hitachi, using Agilent zorbax RX-C18 column (5 μm, 4.6 × 150 mm)). Peptide identity was confirmed using mass spectrometer (Thermo Scientific).

### 2.2 Protein expression and purification

ERK2: ERK2^R65S^ was expressed using a NpT7-5 His_6_-ratERK2 plasmid as described.^25^ After expression, the bacteria expressing the protein were lysed using a microfluidizer. The soluble fraction was separated by centrifugation and loaded on a nickel Sepharose 4 mL column. Elution of the protein was performed using an imidazole gradient, at 100% elution buffer (containing 25 mM Tris-HCl pH = 8.0, 500 mM NaCl, 5% glycerol, 5 mM βME and 300 mM imidazole). The protein was further purified using a size exclusion chromatography Sephacryl S100 500 mL column. Elution was performed using a buffer containing 25 mM Tris-HCl pH = 8.0, 300 mM NaCl, 5% glycerol and 5 mM βME, which was also used as the storage buffer. The kinase purity was confirmed by coomassie staining of SDS-PAGE gel (Fig. S3).

P38γ: For protein expression, a starter of Rosetta cells harboring the expression plasmid (6.5 ml) was grown overnight at 37 °C. Then, the starter was diluted to a final culture volume of 250 ml and further incubated at 37 °Cto O.D. 600 nm = 0.3-0.4 when IPTG was added to a final concentration of 0.3 mM. The culture was further incubated at 30 °Cfor about 5 hours until it reached O.D. 600 nm = 0.5-0.6 and centrifuged at 3,200 rcf at 4 °Cfor 10 minutes. The pellet was suspended in cold sonication buffer (300 mM NaCl, 50 mM Tris-HCl pH = 8.0 and 10 mM imidazole) and re-centrifuged at 2,200 rcf at 4 °Cfor 10 minutes. The pellet was frozen in liquid nitrogen and stored at -80 °C. For protein purification, pellets were thawed on ice and suspended in 10 ml sonication buffer with protease inhibitor cocktail (Protease inhibitor cocktail 1X, Tivan-Biotool). The suspension was sonicated in two identical cycles: 10 seconds sonication, 10 seconds rest (3 repeats), 1 minutes rest, 10 seconds sonication, 10 seconds rest (3 repeats). All sonication cycles were set to an amplitude of 30% and were done on ice. Following sonication, the suspension was centrifuged at 20,000 rcf at 4 °Cfor 40 minutes. The supernatant was loaded into a gravity column containing 1 ml of Ni-NTA beads (PureCube Ni-NTA 40μm 50%, Tivan) that were prewashed with 10 ml of cold sonication buffer. The flow-through was collected, discarded and the beads were re-washed with 20 ml of cold sonication buffer that was collected and discarded. Finally, the beads were incubated in 3 ml of elution buffer (300 mM NaCl, 50 mM Tris-HCl pH = 8.0 and 250 mM Imidazole) for 5 minutes and three elution fractions of 1 ml each were collected. These fractions were then dialyzed in dialysis membranes (Medicell Membranes Ltd Molecular weight Cut Off 12-14000 Daltons) against dialysis buffer (12.5 mM HEPES pH = 7.5, 200 mM KCl, 0.5 mM DTT and 0.625% glycerol) in gentle stirring at 4 °Covernight. Proteins were collected and stored in small working aliquots (15-100 µl) at -80 °Cafter freezing in liquid nitrogen.

### 2.3 Fluorescence anisotropy

All anisotropy measurements were performed with fluorescein-labeled ELK-1_312-321_ peptide (**2**) (FL-GGGWKGRKPRDLGL-NH_2_), using a spectrofluorimeter (Perkin-Elmer Life Sciences LS-55 spectrofluorimeter equipped with a Hamilton microlab M dispenser). The peptide (**2**) (1 ml, 100 nM) was dissolved in 25 mM tris buffer, pH 8, 100 mM NaCl, 10% glycerol, and 5 mM β-mercaptoethanol). FL-ELK-1_312-321_ (1 ml, 100 nM) was placed in the cuvette and the appropriate ERK2 construct (250 µL, 200 µM) was placed in the dispenser. The aliquots (10-20 µL) were added at 1.5 min intervals followed by the solution were then stirred for 30 s, and its fluorescence anisotropy was measured at 10 °Con excitation at 492 nm and emission at 520 nm. The data was analyzed using Origin 8 (OriginLab) and fit to a 1:1 binding model Eq. (1).

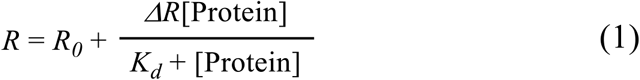

Where *R*: measured fluorescence anisotropy, *R*_*0*_: anisotropy value corresponding to the free peptide, *⊿R*: change in the amplitude of the fluorescence anisotropy, Protein: protein concentration, and *K*_*d*_: dissociation constant.

### 2.4 Electrode preparation

Initially, gold electrodes were polished with micro-cloth pads (Buehler, Lake Bluff, IL) with two separate alumina suspension (Buehler) of 1 μM, and 0.05 μM particle sizes, and cleaned with sonication in ethanol:TDW (1:1) for 10 minutes and subsequently washed with TDW. The cleaned electrodes were immediately modified by drop casting with 100 µM solution of ELK-1_312-321_ (**1**) peptide in ammonium acetate buffer (pH 6.8) at 25 °Cin the incubator for 16 h, resulting in self-assembled peptide monolayer. The peptide (**1**) was coupled through cysteine from N-terminus and three glycine as linker (Fig. 2). The adsorption of peptide on gold surface was monitored by electrochemical analysis. This chemisorption resulted in enhancement of resistance, confirmed the final electrode preparation.

### 2.5 Electrochemical measurements

Electrochemical impedance spectroscopy (EIS) analysis was conducted with Bio-Logic SP-300 potentiostat (Bio-Logic Science Instruments, France), assisted by EC-LAB software package. The electrochemical measurements were carried out in a regular three-electrode electrochemical cell at 25 °C. The reference electrode (Ag/AgCl), Pt rode as counter electrode and gold disc (diameter of 2 mm) as a working electrode (Au). The measurements were carried at first with bare gold electrode followed by peptide immobilization and then post incubation with kinase. EIS scans were done in EIS solution of 5.0 mM K_3_Fe(CN)_6_, 5.0 mM K_4_Fe(CN)_6_ (RedOx species), and 0.1 M KCl as supporting electrolyte. The interaction of the ERK2 with surface bound ELK-1 peptide (**1**) was also performed at various concentrations (0.005 μM, 0.05 μM, 0.25 μM, 0.5 μM, 1.0 μM, 2.0 μM, 4.0 μM and 8.0 μM), using an electrochemical impedance technique in 50 mM ammonium acetate and 100 mM KCl buffer solution. All the impedance data were fitted with the circuit R_S_(R_CT_|W)||Q, where Q is a constant phase element, which describes a non-ideal capacitor.

### 2.6 Surface characterization

The gold surfaces were prepared by Au (100 nm layer) evaporation on the top of Cr layer (10 nm), which evaporated on the top of n-type Si wafer ⟨100⟩. XPS analysis was done using a monochromatic Al Kα X-ray source on a Kratos AXIS ULTRA instrument. The XPS was used for elemental analysis of the organic layer. Atomic force microscopy (AFM, Bruker) was acquired in tapping mode to monitor topography homogeneity of the layer. Polarization modulation infrared reflection absorption spectroscopy (PM-IRRAS) measurements were conducted at room temperature under positive nitrogen gas pressure on a reflection-absorption cell (Harrick, Inc.) with a PM-FTIR spectrometer (PMA-50 coupled to Vertex V70, Bruker). The signal was collected from modified Au surfaces by 2048 scans with a resolution of 4 cm^-1^ using a mercury cadmium telluride detector.

## 3. Results and discussion

### 3.1 Peptide design

To design a peptide-based sensor for ERK2 that is based on an interaction rather than the catalytic activity, we looked for sequences from ERK2-binding proteins that bind the docking site. We selected a peptide derived the docking motif of ELK-1,^23^ one of the substrate proteins ofERK2. ELK-1 is a transcription factor that regulates immediate early gene (IEG) expression through the serum response element (SRE)^26^ DNA consensus site and its phosphorylation occurs in response to MAPKs and ERKs. (ELK-1 312-321, ^312^KGRKPRDLEL^321^).^27^ We added a cysteine residue for binding the gold electrode (Fig. S1) and a tryptophan residue at the N-terminus of the peptide for concentration measurements using UV/Vis spectroscopy. We also added a three glycine linker at the N-terminus. The resulting sequence of the peptide was CGGGWKGRKPRDLEL. The peptide was also synthesized with fluorescein at N-terminus for fluorescence anisotropy binding studies (Table S1).

The interaction between ERK2 and the ELK-1_312-321_ peptide (**2**) was quantified using fluorescence anisotropy and the *K*_*d*_ was found to be 14.7 (±0.2) µM (Fig. S2).

### 3.2 Construction of electrochemical peptide-based biosensor

Peptide (**1**) was immobilized on the gold electrode surface in order to assemble the peptidic monolayer. The cysteine at N-terminus mediates strong chemisorption of peptide to the gold electrode.^28^ Subsequently, the electrodes were incubated with ERK2 by drop casting for 1 h at 25 °Cin ammonium acetate buffer (Fig. 2). EIS measurements were used to monitor each step of the electrode interfacial modifications, and the detection of the interaction between peptide (**1**) and ERK2.

Next, we tested whether the peptide-bound gold electrode can sense ERK2 (Fig. 3). Upon exposure to ERK2, the impedance increased from 834±166 Ω to 1456±291 Ω. The R_CT_ value of electrode with ERK2 increased by 70-80%, whereas it remained ∼10% within the error range with the buffer only (Fig. S4). The error could be associated with the impedance drift due to partial coverage of peptide monolayer.^29, 30^ This may indicate binding of ERK2 to the peptide monolayer and not rearengment in the monolayer packing.

**Fig. 3.**
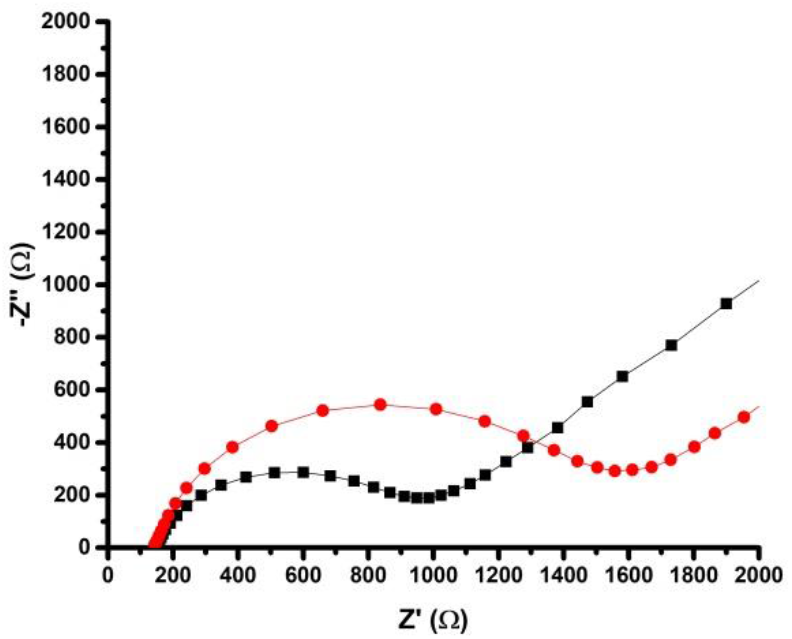
Sensing ERK2 by the ELK-1_312-321_ electrode. Black square - the R_CT_ of the peptide monolayer (R_CT_ = 834±166 Ω). Red circle - the R_CT_ of the monolayer following interaction with ERK2 (R_CT_ = 1456±291 Ω).

### 3.3 Dose dependence of ERK2 sensing

To reveal the sensitivity of the biosensor we carried out a dose response study. The electrodes with the assembled peptide monolayer were exposed to various concentrations of ERK2 0.005 μM, 0.05 μM, 0.25 μM, 0.5 μM, 1.0 μM, 2.0 μM, 4.0 μM, and 8.0 μM and the changes in the R_CT_ values are shown in Fig. 4a. The results show that the sensor was sensitive towards ERK2 from 0.25 μM and above. The lowest possible concentration for the ERK2 was calculated as 0.52±0.08 μM. The measured lowest possible concentration is at the same order of magnitude of the measured intracellular concentration (0.8 μM) of ERK2.^31^ The amplification of ERK2 occurs in the different cancers such as ovarian, lung, bladder, and breast cancers by 12% to 21%.^32^ So our assay is capable to measure the ERK2 as a cancer biomarker.

**Fig. 4.**
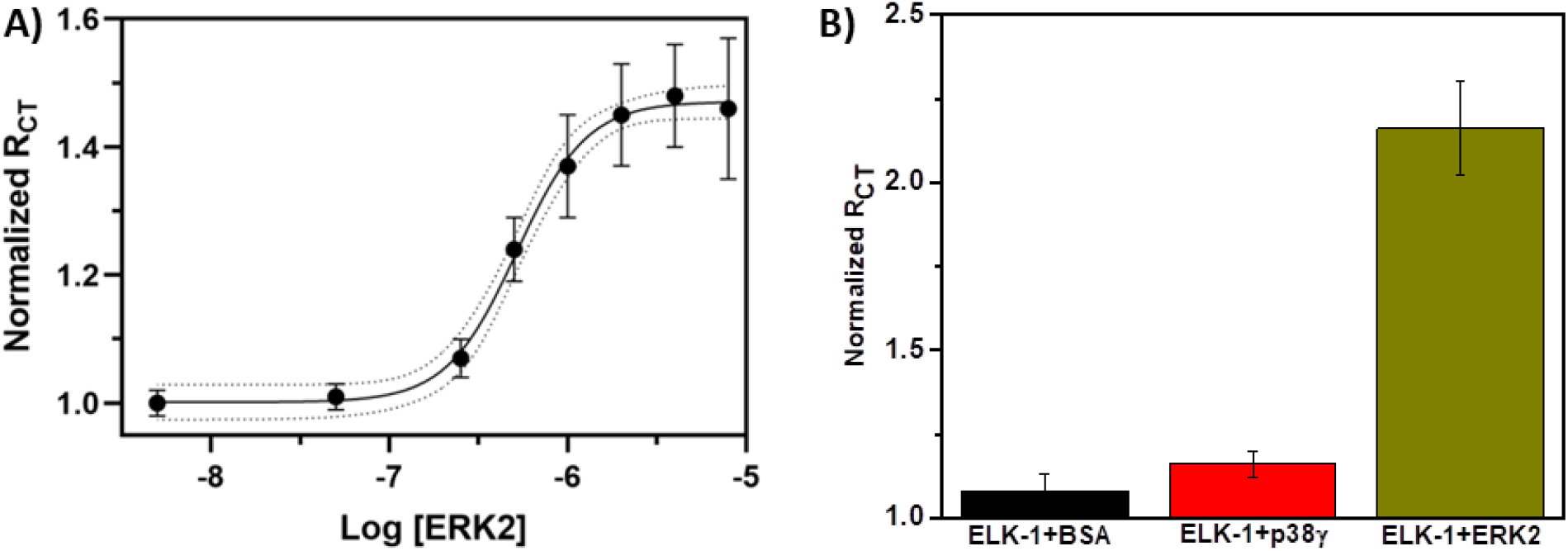
A) EIS-derived dose response of peptide biosensor to ERK2. Shown are normalized R_CT_ values after incubation with different concentrations of ERK2 solutions. Y-axis normalized R_CT_ is calculated by the final R_CT_ value divided by the initial R_CT_. B) The specificity of the peptidic biosensors towards ERK2. The EIS experiments show that after exposure to BSA and p38γ, the changes in the properties of the layers were not significant in comparison to electrodes exposed to ERK2. The shown R_CT_ values after incubation with BSA, p38γ and ERK2 are normalized by initial values.

### 3.4 The peptide biosensors are selective towards ERK2

To determine whether the sensing of ERK2 is selective, we carried out the electrochemical impedance measurements with BSA and p38γ (Fig. 4b). BSA is a highly abundant protein while p38γ is a kinase from the MAPK family, to which ERK2 also belongs.^27^ The results show no significant changes in R_CT_ values for the electrodes following exposure to BSA and p38γ, demonstrating the selectivity of the peptidic biosensor for ERK2. The selectivity between the kinases from the same family is highly significant, showing the strong effect and high selectivity of the docking site interaction.

### 3.5 Surface analysis demonstrates the thickness measurement for peptide-kinase bilayer

To further understand the mechanism of ERK2 sensing, we measured the thickness of the peptide monolayer by variable angle spectroscopic ellipsometry (VASE). The peptide (**1**) was immobilized on the gold surface and observed thickness was 3.6±0.1 nm. The thickness of the peptide monolayer increased by 2.5±0.05 nm following the exposure to ERK2, indicating that ERK2 is indeed present on the peptide monolayer, see Table S2. The increase in the optical thickness by 2.4 nm supports the added layer of ERK2 on the peptide monolayer. The dimensions of the ERK2 compared to the peptide in this range over rules the conformational changes of peptide monolayer.

Next, we monitored the topography of ELK-1_312-321_ peptide monolayer on ultra-flat gold substrate^33^ after adsorption and after exposure to ERK2 using AFM. The roughness of the ultra-flat gold substrate was analyzed before peptide immobilization, resulting in RMS roughness of 2.1±0.4 Å (Fig. 5A). AFM scans of the gold substrate after peptide monolayer formation resulted in 2.6±0.6 Å (Fig. 5B). After incubation with ERK2 the roughness substantially increased to 5.2±1.4 Å (Fig. 5C). These results show a considerable change in the average roughness, which clearly indicates the binding of ERK2 to the peptide layer.

**Fig. 5.**
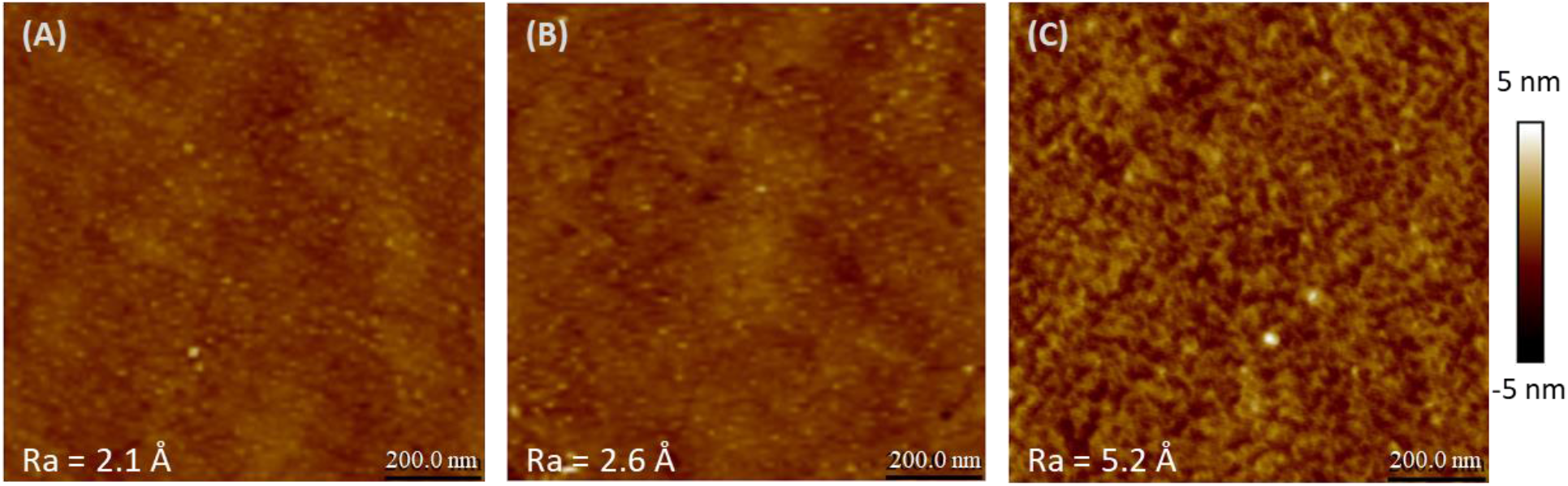
AFM topography images for peptide **1**-assembled monolayer on ultra-flat gold substrates and following ERK2 incubation: (A) bare gold substrate; (B) peptide ELK-1_312-321_ assembled monolayer; (C) after 1 h incubation with ERK2. Surface roughness for all samples was determined on a 1.0×1.0 μm scanned area.

To characterize the assembled peptide monolayer on the surface we carried out XPS analysis. The gold substrates with the peptide absorbed showed a characteristic peak at binding energies of 400.3 and 161.5 eV. N 1s and S 2p are related to these two peaks (Fig. S5 and S6, see ESI). The bare Au was scanned as a reference and shows absence of the peaks related to the N and S. This confirms the adsorption of peptide on the Au substrates. However, analysis of bare Au represents only C 1s-related peak at 283.8 eV (Fig. S7, see ESI), which may be the presence of carbon impurities. Analysis of Au-ELK-1_312-321_ peptide monolayer surfaces (Fig. S7, see ESI) represents the C 1s peaks at 285 and 286.5 eV. The Au substrates with ERK2 incubates on Au-ELK-1_312-321_ peptide monolayer surfaces, shows C 1s peaks at 285.3, 286.6, and 288.3 eV. These peaks are indicated as C-C, C-N/C-O, and O=C-N respectively and consequently provides evidence for the peptide functionalities on the gold surface.^34^ The increase in the percentage of N, S and C upon incubation with ERK2 indicates the interaction between ELK-1_312-321_ and ERK2 (Table S3). The substantial increase in S 2p signal indicates the more number of cysteines coming from ERK2. Along with decrease in the Au percentage makes strong point for added layer of the ERK2. Moreover, we determined the thickness of these substrates by XPS. All the XPS analyses were performed with two substrates to avoid any artifact. The thickness of the substrates increased after exposure to ERK2 (Table S4). The difference in the thickness of layer was ∼ 2.28 nm, supporting the binding of ERK2 to the peptide monolayer and in line with the ellipsometric optical thickness measurements.

We used polarization modulation IR reflection-absorption spectroscopy (PM-IRRAS) measurements to monitor the adsorbed layer of ERK2 on the ELK-1_312-321_ peptide monolayer. We focused mostly on changes of signals in the amide I and amide II regions. The amide I band (1600-1700 cm^−1^) is associated to the C=O stretching frequency of the amide groups and to in-plane N-H bending. The amide II band (1480-1575 cm^−1^) refers mainly to the in-plane N-H bending and from the C-N stretching.^35, 36^ Significant change was observed in the absorbance at 1542 cm^-1^ in the amide II region upon exposure of the peptide monolayer to ERK2 (Fig. 6). No significant difference was observed in the amide I region. The change that was related to the amide II region (1416-1593 cm^-1^) clearly validates the additional C-N stretching and N-H bending from ERK2 in comparison to ELK-1_312-321_. Also, it indicates the presence of ERK2 on the surface. Other than the amide I and amide II regions there are spikes in the spectrum (1000-1420 cm^-1^) showing the presence of many functional groups, originating from the adsorbed ERK2 layer.

**Fig. 6.**
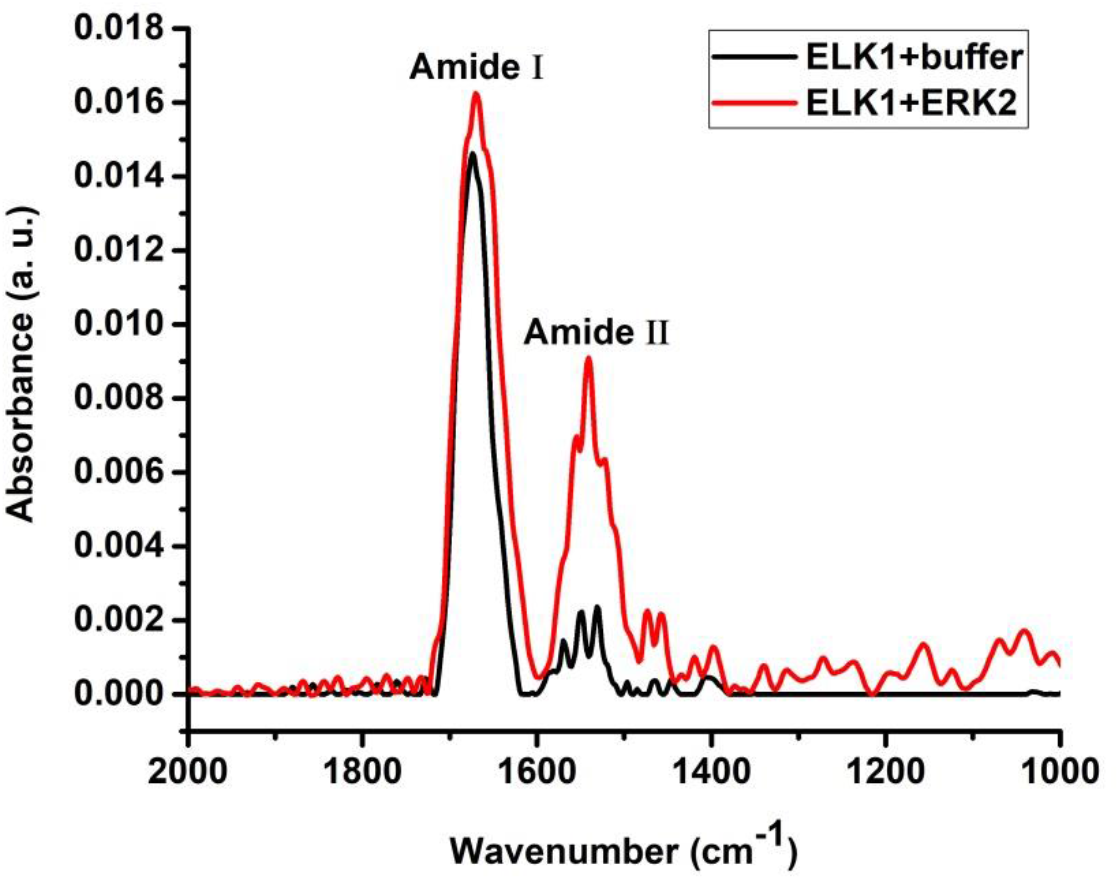
PM-IRRAS scan: black - ELK-1_312-321_ peptide monolayer; red - ELK-1_312-321_ peptide monolayer after incubation with ERK2 (1 µM) for 1 h at 25 °C.

Here we present a novel approach for the development of a kinase biosensor based on the docking-site interactions of the kinase with a peptide derived from its partner protein. This approach does not require catalytic activity for sensing. The peptide from D domain of ELK-1, which binds with ERK2 at the DRS with micromolar affinity,^20^ was utilized as a surface bound recognition layer.

### 3.6 Interactions – based sensing vs. catalytic site based sensing

The tools available so far for the detection of enzymes are based on sensing enzymatic activities such as identifying the products of enzymatic reactions,^18^ post-translational modifications,^37^ chemiluminescence^3^ and similar methods. These methods rely on the particular enzymatic activities and their sensitivity is related to the extent and specificity of the catalytic activity. Kinases are sensed based on their catalytic active sites and those may not be very specific for relevant substrates.^38^ For example, casein kinase (CK2) phosphorylates at least 160 proteins, making recognition through its catalytic site non-specific. ERK2 also phosphorylates many substrate proteins.^21^ Thus, developing biosensors based on the catalytic site is likely to result in lack of sufficient selectivity, which could lead to false positive signals from other kinases. The lack of substrate specificity of kinases *e.i*., reaction rate based selectivity^15^ led us to explore alternatives for sensing kinases by targeting other sites and functionalities. The approach presented herein, based on the interactions of the DRS, can offer a high level of selectivity. The biosenors demonstrate selectivity for sensing ERK2 and not the kinase from the same family p38γ (Fig. 4b), indicating the usefulness of using docking-site interactions over using the catalytic site based sensing for achieveing selectivity.

### 3.7 The advantages of electrochemical sensing compared to ELISA

ELISA is the common method that is widely used in the clinical diagnosis and target identification via antibody specificity. It offers cheap, easy and accurate detection and quantification of biological targets.

Using electrochemical methods has several advantages over using ELISA: In ELISA, primary and secondary antibodies have to be used. These antibodies are expensive to produce, should be stored under conditions that retain their structure and activity and should not lose their activity upon immobilization if required.^7^ The need to detect pathogenic species at harsh conditions further limits the shelf life of antibody functionalized sensors.^39^ Enzymes sandwiched between antibodies were shown to function as electrochemical biosensors through gold electrodes fabricated with microporous nylon membranes,^40^ but this results in complex systems using enzyme linked antibodies. The antibodies can be expensive and have stability issues. Our docking-site sensor, which is peptide-based, does not require enzyme activity paired with antibodies, but rather requires the synthesis of simple and stable peptides and not of unstable and expensive antibodies. Another advantage of the approach presented herein is that it is label free and simple in terms of catalytic activity choices.^41^ The technique is cheap, selective and is simple to operate and analyze. The electrochemical sensing assay with BSA showed selectivity towards ERK2 (Fig. 4b). The measured lowest possible concentration (0.52±0.08 μM) from the dose response study is close to the cellular level (0.8 μM) of the kinase concentration,^31^ which makes the system practical for sensing kinases from in vivo samples.

## 4. Conclusion

In summary, the sensor presented herein can detect enzymes without the requirement for specific antibodies or electrochemical labels. The selectivity between the kinases found to be crucial. It detects the ERK2 at its cellular level brings in the realistic use in advancement of biosensors. Docking site based detection of kinases could opens up new spectrum to detect kinases precisely as compare to the enzymatic acitivity. The approach could be applied for example for sensing cancer-related enzymes and thus detect early onset of cancer. This could be later applied for other disease-related enzymes as well.

## Acknowledgments

AF and SY thank the support of the Israel Science Foundation (ISF) grant number 1628/18. This work was supported by the Academia Sinica and the Hebrew University of Jerusalem joint research program in Nanoscience and Nanotechnology and The Minerva Center for Bio-Hybrid complex systems. We thanks to Prof. D. Engelberg from the Hebrew University for contributing the p38γ kinase. S.Y. thanks the Binjamin H. Birstein Chair in Chemistry. AF thanks the Saerree K. and Louis P. Fiedler Chair in Chemistry.

