## Supplemental file for "Electrochemical biosensors based on peptide-kinase interactions at the kinase docking site"

### 1. Additional results and data

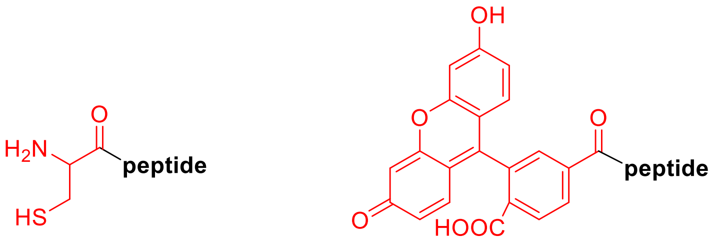


**1** **2**

**Peptide**-Gly-Gly-Gly-Trp-Lys-Gly-Arg-Lys-Pro-Arg-Asp-Leu-Glu-Leu

**Fig. S1.** Synthesized peptides for EIS and fluorescence anisotropy

Table S1. Peptide sequences used in this study

| **Entry** | **Sequence** |
|---|---|
| 1 | CGGGWKGRKPRDLEL (**1**) |
| 2 | FL-GGGWKGRKPRDLEL (**2**) |

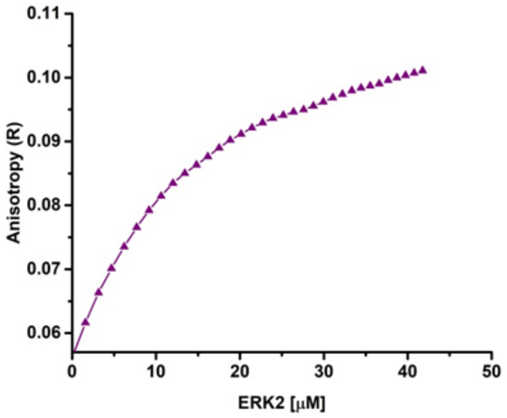


**Fig. S2**. Fluorescence anisotropy binding studies of ERK2 and FL-ELK-$1_{312\text{-}321}$ (**2**). $K_d$ was found to be 14.7±0.2 μM.

(A) (B)

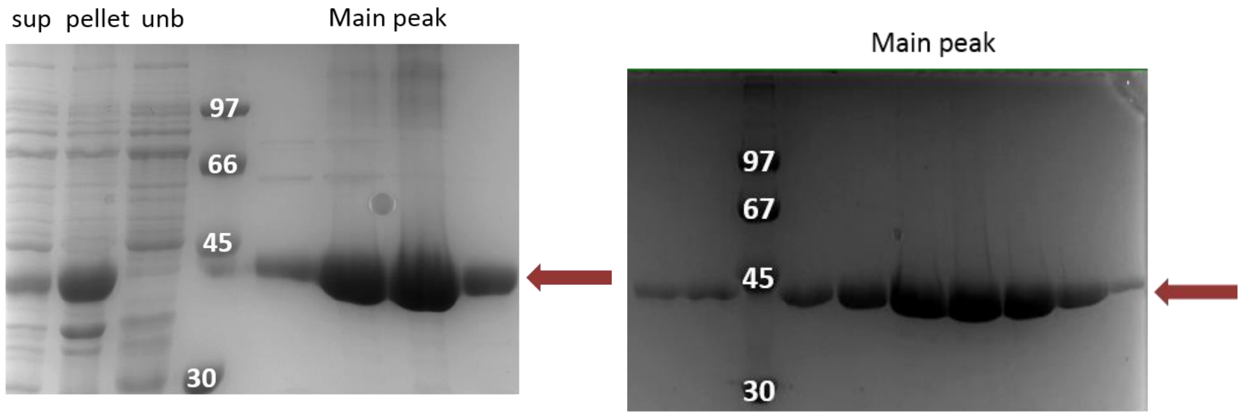


**Fig. S3**. Purification of ERK2$^{R65S}$ -(A) Coomassie staining of a SDS-PAGE gel of ERK$^{R65S}$ after the bacteria lysis and nickel affinity chromatography column; (B) Coomassie staining of a SDS-PAGE gel of ERK$^{R65S}$ after the size-exclusion chromatography column.

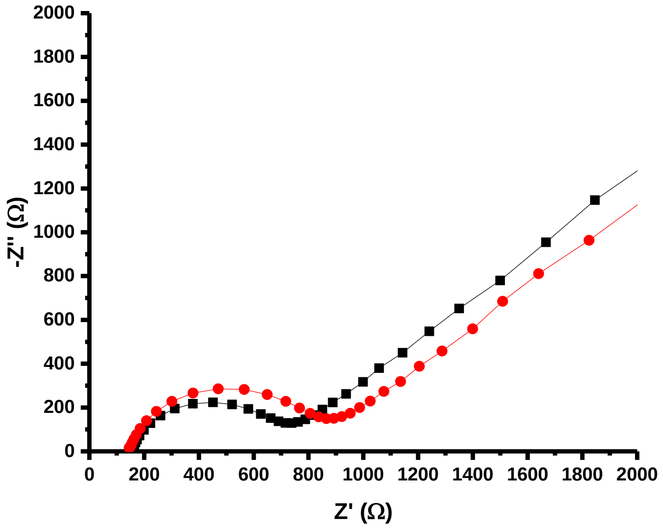


**Fig. S4.** Control experiment between ELK-1$_{312\text{-}321}$-buffer. Black square - the $R_{CT}$ of the peptide monolayer ($R_{CT}$ = 610±122 Ω). Red circle - the $R_{CT}$ of the monolayer after buffer addition ($R_{CT}$ = 728±144 Ω).

**Table S2.** The thickness increment is ~ 2.5 nm by ellipsometry.

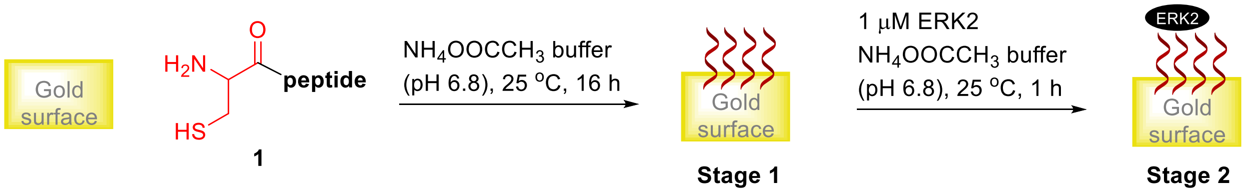


| Entry | Substrates | Thickness (nm) |
|---|---|---|
| 1 | ELK-$1_{312\text{-}321}$ | $3.5^a \pm 0.1$ |
| 2 | ELK-$1_{312\text{-}321}$ + ERK2 | $6.0^a \pm 0.6$ |

[a] The mean of three VASE values from the substrate.

Representative data of VASE for one value of each entry:

A) Entry 1

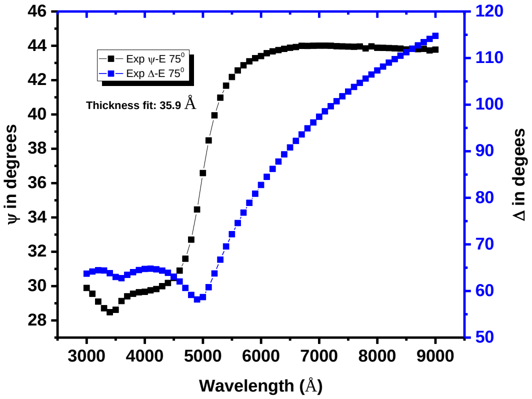


B) Entry 2

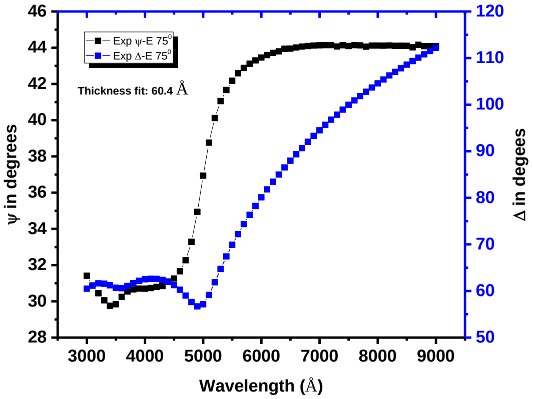

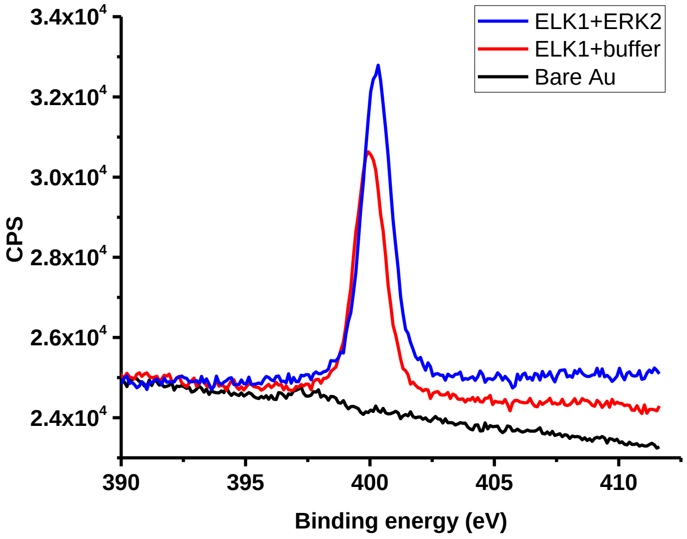


**Fig. S5.** XPS analysis for N 1s: Black - bare Au; red - ELK-1+buffer and blue - ELK-1+ERK2. Binding energy peaks at 399.9 and 400.3 eV respectively.

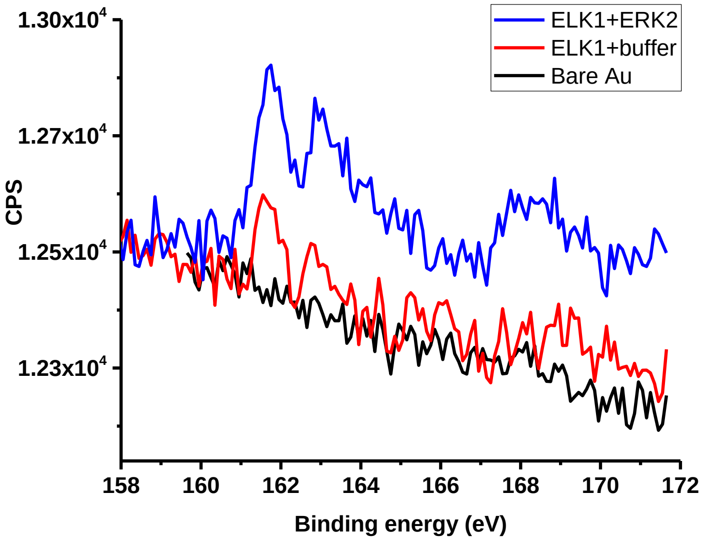


**Fig. S6.** XPS analysis for S 2p: Black - bare Au; red - ELK-1+buffer and blue - ELK-1+ERK2. Binding energy peaks at 161.5, and 161.7 respectively.

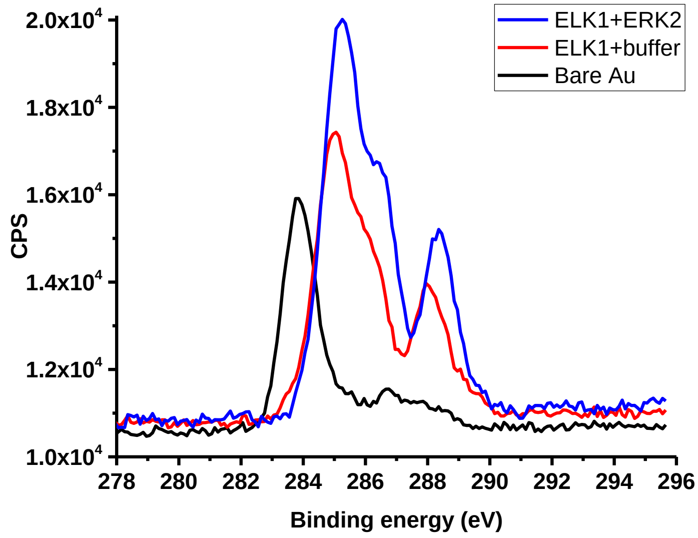


**Fig. S7.** XPS analysis for C 1s (C-H): Black - bare Au; red - ELK-1+buffer and blue - ELK-1+ERK2.

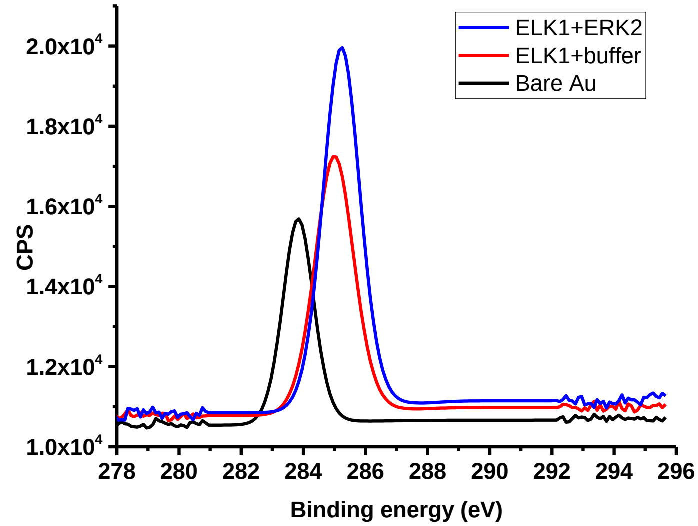


**Fig. S8.** XPS analysis for C 1s (C-O): Black - bare Au; red - ELK-1+buffer and blue - ELK-1+ERK2. Binding energy peaks at 283.8, 285, and 285.3 eV respectively.

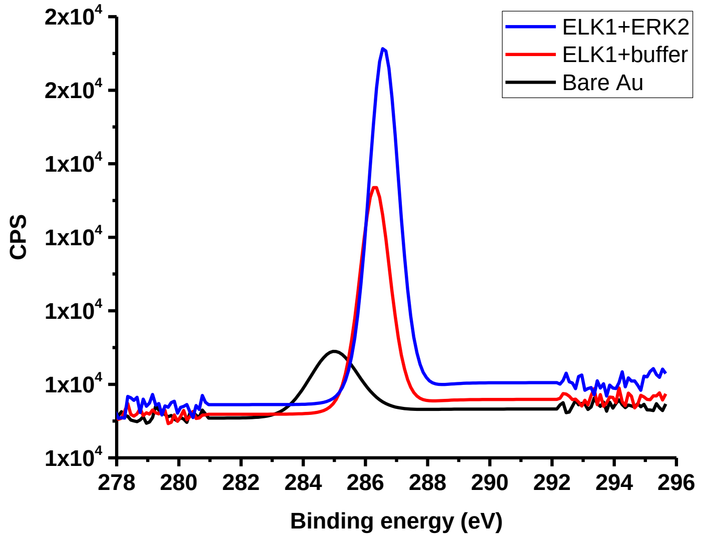


**Fig. S9.** XPS analysis for C 1s (C=O): Black - bare Au; red - ELK-1+buffer and blue - ELK-1+ERK2. Binding energy peaks at 285, 286.3, and 286.6 eV respectively.

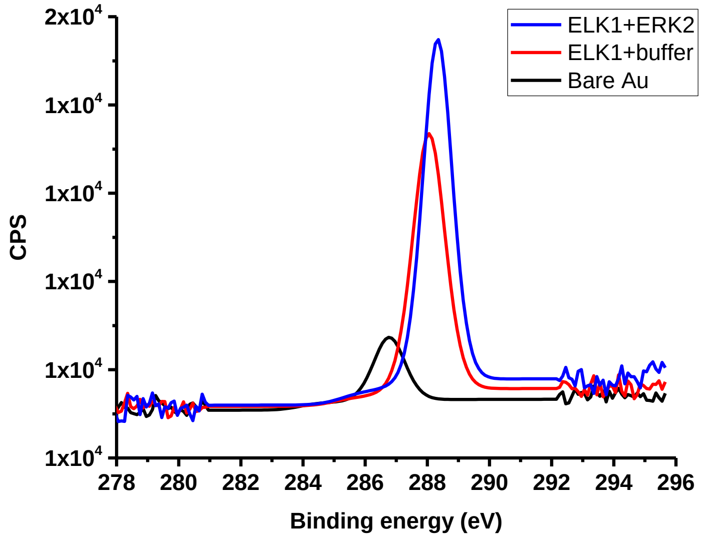


**Fig. S10.** XPS analysis for C 1s (COOH): Black - bare Au; red - ELK-1+buffer and blue - ELK-1+ERK2. Binding energy peaks at 286.7, 288, and 288.3eV respectively.

**Table S3**. XPS analysis of gold substrates

| Entry | Atom | Atomic concentration [%][a] | | |
|---|---|---|---|---|
| | | **ELK-1+buffer** | **ELK-1+ERK2** | **Bare Au** |
| 1 | Au 4f | 1 | 1 | 1 |
| 2 | S 2p | 0.006 | 0.08 | 0 |
| 3 | C 1s | 1.7 | 4.8 | 0.7 |
| 4 | N 1s | 0.5 | 1.4 | 0.02 |
| 5 | O 1s | 0.5 | 1.4 | 0.2 |

[a] The atomic concentration are normalized values after incubation with buffer and ERK2 solutions. The normalized atomic concentration is calculated by the atomic concentration value divided by the percentage of Au.

**Table S4**. Thickness measurement by XPS

| Entry | Substrates | Thickness (nm) |
|---|---|---|
| 1 | ELK-$1_{312\text{-}321}$ | 2.75±0.03 |
| 2 | ELK-$1_{312\text{-}321}$ + ERK2 | 5.03±0.07 |
